# Biofilms Confer Resistance to Simulated Extra-terrestrial Geochemical Extremes

**DOI:** 10.1101/099374

**Authors:** Adam H. Stevens, Delma Childers, Mark Fox-Powell, Charles S. Cockell

## Abstract

Biofilms improve microbes’ resistance to a variety of extreme physical and chemical conditions on Earth. The discovery of putative aqueous environments on other planetary bodies such as Mars motivates an interest in understanding the viability of life, and the potential role of biofilms, in previously unexplored geochemical extremes. We investigated the loss of viability of planktonic cells and biofilms of *Sphingomonas desiccabilis* (a Gram-negative, desiccation resistant, soil crust-forming organism) to simulated Martian brines. These brines were produced from geochemical modelling of past aqueous environments on Mars, and their high sulfate concentrations make them different to most terrestrial brines, although similar briny environments have been found in locations such as the Basque Lakes in Canada or in deep subsurface groundwater systems. Biofilms grown on basaltic scoria were subjected to the simulated martian brines and the viability of cells was measured over time and compared to equivalent planktonic cultures. Crystal violet assay was used to measure how the biomass of the biofilms changed over time in response to the brines. While certain brines were highly hostile to microbial viability, we found that biofilms that were desiccated prior to being treated with brines maintained viability over a longer treatment period when compared to planktonic cells. Our results show that biofilms confer short-term protection to the harsh osmotic, ionic, and acidic conditions of Mars-relevant brines. However, in the most extreme simulated brines, even biofilms eventually lost viability. By demonstrating that biofilms confer protection to conditions that are potentially analogous to current day recurrent slope lineae (thought to be produced by the flow of briny fluids) on Mars, our results show that contaminant biofilm-forming microorganisms may have a greater chance of surviving in so-called ‘Special Regions’ on Mars, with implications for planetary protection in missions that aim to explore these regions.

## 2 Introduction

Biofilms are produced by a variety of organisms and generally consist of cells attached to a surface (and each other) in an exuded matrix of extracellular polymeric substance (EPS) [1]. Biofilms can be formed by single species or complex microbial communities and are generally produced in response to the presence of a surface or external cues and stressors. Microbial biofilms increase the resistance of the inhabitants to a number of extreme conditions including immunological aggression [2], antibiotics [3], bactericidal chemicals [4], rapid changes in temperature and pH [5], UV radiation [6] and salinity [7,8]. Biofilm-producing microorganisms are therefore present and thrive in a range of environments on Earth and may have been involved at an early stage as one of the steps in the development of the first cells [9].

Although biofilms confer resistance to extremes found on Earth, their ability to provide protection against entirely novel chemical conditions found on other planetary bodies remains unknown. The search for extra-terrestrial habitable environments centres on the availability of liquid water, and aqueous environments are now known to exist on a number of planetary bodies including Mars, Europa, Enceladus and Titan [10–14]. Many of these aqueous environments contain combinations of anions that are rare on Earth. For example, the high concentrations of sulfate and other divalent ions in Martian brines, a consequence of the specific geological history of that planet, have consequences for habitability, making ionic strength an important determinant of whether the brines will support life [15]. Recent observations suggest that recurrent slope lineae (RSL) features on Mars are created by the action of flowing water which is saturated with dissolved perchlorate salts [16]. Such brine compositions have not been found on Earth. In this study, we investigated the resistance of biofilms to aqueous environments analogous to environments on Mars such as RSL currently potentially active [16] or in quasi-tidal areas that existed during the phase of the planet’s history when it lost its surface water to atmospheric escape or to retreat into subsurface ice [17,18], but also have implications for the icy moons in the outer solar system, which are thought to have high levels of dissolved salts in their liquid water interiors [19].

We hypothesise that production of biofilms might protect microorganisms and allow them to maintain viability following exposure to compositionally diverse brines with high levels of dissolved salts. Our results have implications for the adaptations that would be required by any putative life in extra-terrestrial environments and for planetary protection, *i.e.* whether terrestrial organisms accidently introduced into extra-terrestrial brines could persist.

## 3 Results

### 3.1 Viability after brining

Figures 1–4 show the viability of *S. desiccabilis* biofilms over time when immersed in Mars-analogue brines. Each time point was an independent culture (triplicate pieces of scoria with biofilm on them) exposed to brine for a given length of time. Experimental replicates are collected on the same axes. Error bars are the standard error of the biological triplicates from each experiment. Best-fit lines were calculated using a linear regression of all experimental replicates.

In Type I brines (Fig. 1) there was only minor decreases in viability over more than 300 minutes when dried or when hydrated. In some cases there appears to be growth during brine immersion (Ib). In brine IIa (Fig. 2a), viability was maintained for over 300 minutes whether the biofilms had been dried or not. In brine IIb (Fig 2b) the dried biofilms showed a decrease in viability, decreasing to below detection levels after around 240 minutes. Hydrated biofilms showed a more rapid decrease in viability, as viable cells were unrecoverable after less than 30 minutes of brine treatment. In type III brines (Fig. 3), viability of cells in the hydrated biofilms reduced to below detection limits after around 30 minutes, which was the shortest timepoint measured. There was more variation in the maintenance of viability of dessicated biofilms treated with the type III brines. For brine IIIa there was a reduction in cell viability over the time course, but not to below detection limits, and not in a linear fashion. In brine IIIb there was a rapid loss of viability to below detection levels within approximately the first 60 minutes, though individual cultures in some experimental replicates showed some measured viability at longer timescales. Brine IIIb has the lowest water activity, the lowest pH and highest ionic strength of all the brines tested. There was a rapid and complete loss of viability in around 100 minutes with both desiccated and hydrated biofilms in type IV brine (Figure 4).

**Figure 1.**
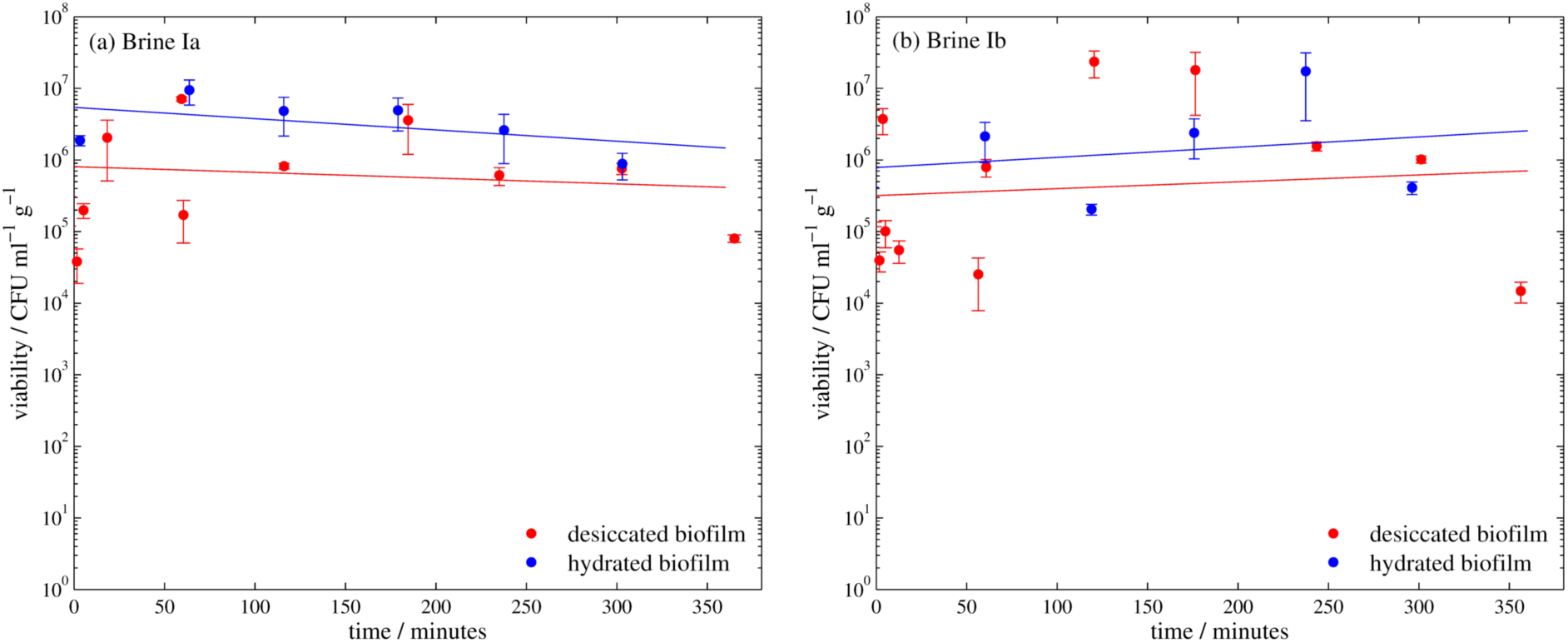
*Viability of S.* desiccabilis *cells in desiccated and hydrated biofilms after periods of immersion in Type I brines, measured by counting colony forming units (CFU). Each data point is a biological triplicate result, with standard error bars, and trend lines are plotted as linear regressions of data from several systematic replicates.*

**Figure 2.**
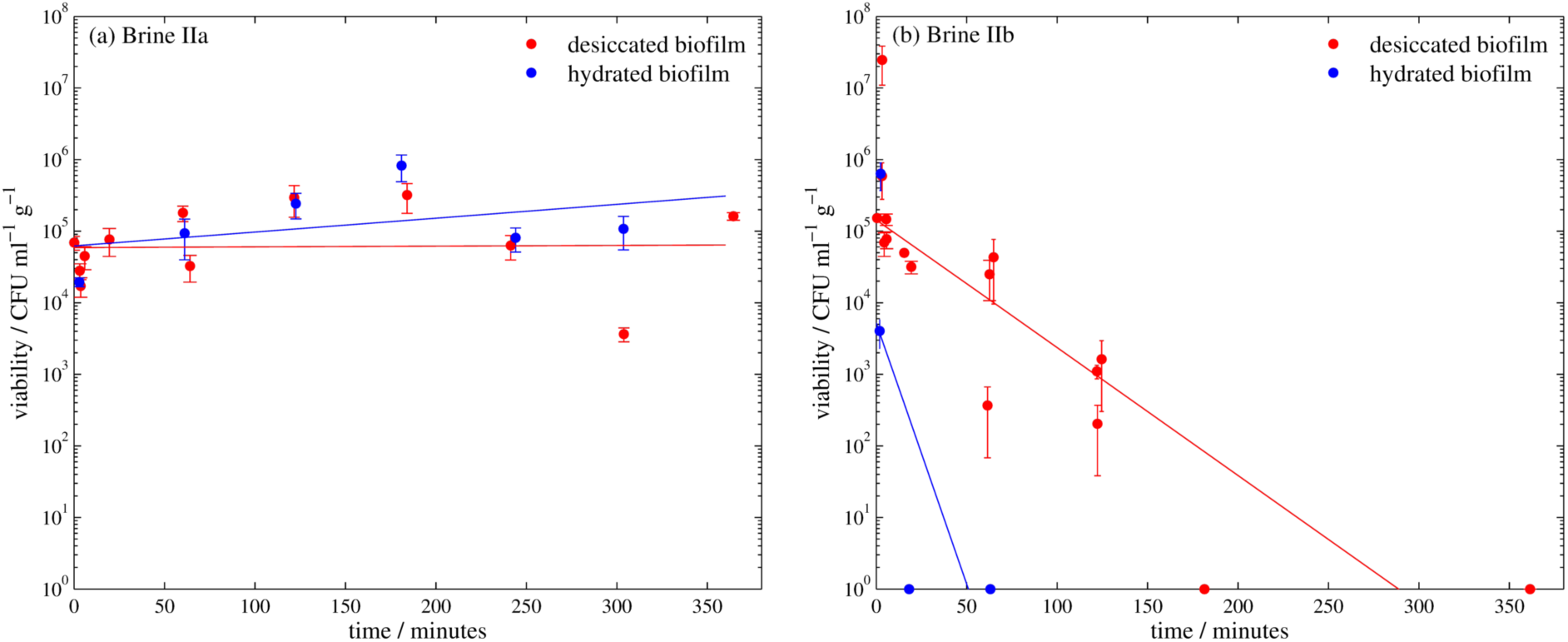
*Viability of S.* desiccabilis *cells in desiccated and hydrated biofilms after periods of immersion in Type II brines, measured by counting colony forming units (CFU). Where no CFUs were countable, results are plotted at 10^0^. Each data point is a biological triplicate result, with standard error bars, and trend lines are plotted as linear regressions of data from several systematic replicates.*

**Figure 3.**
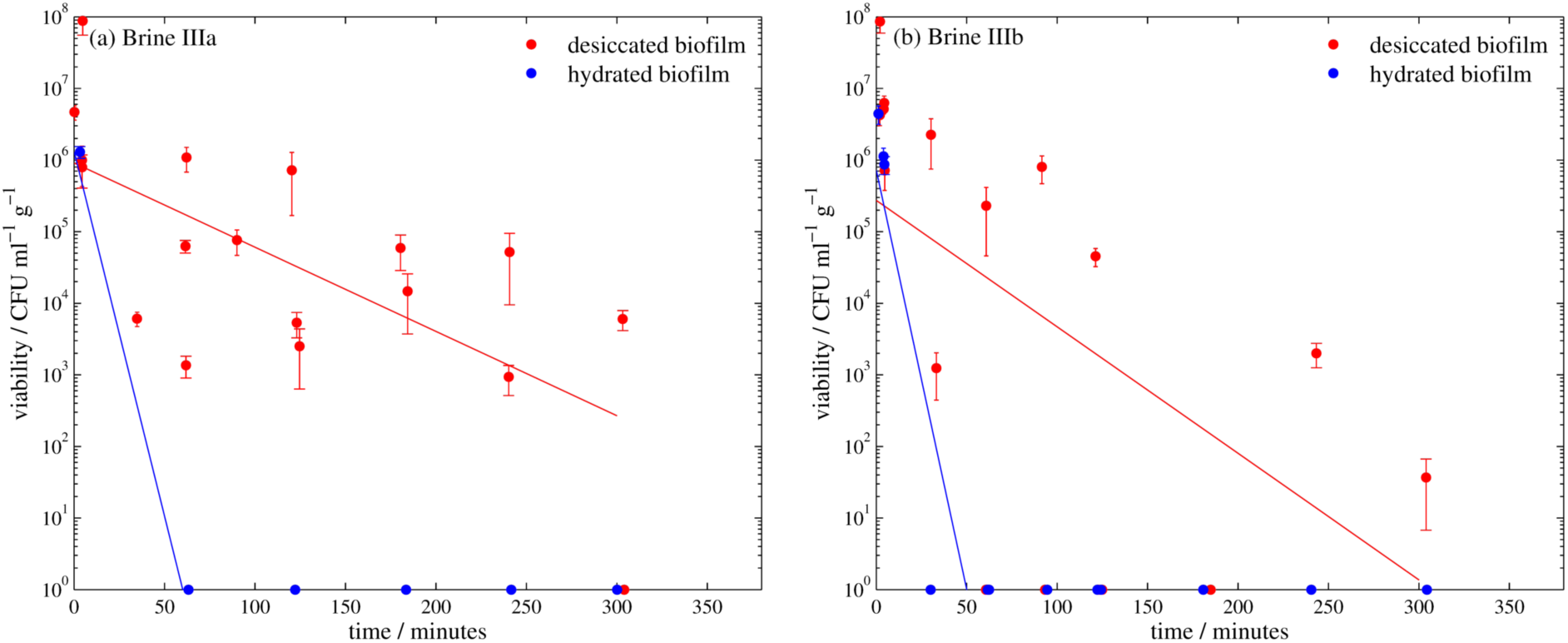
*Viability of S.* desiccabilis *cells in desiccated and hydrated biofilms after periods of immersion in Type III brines, measured by counting colony forming units (CFU). Where no CFUs were countable, results are plotted at 10^0^. Each data point is a biological triplicate result, with standard error bars, and trend lines are plotted as linear regressions of data from several systematic replicates.*

**Figure 4.**
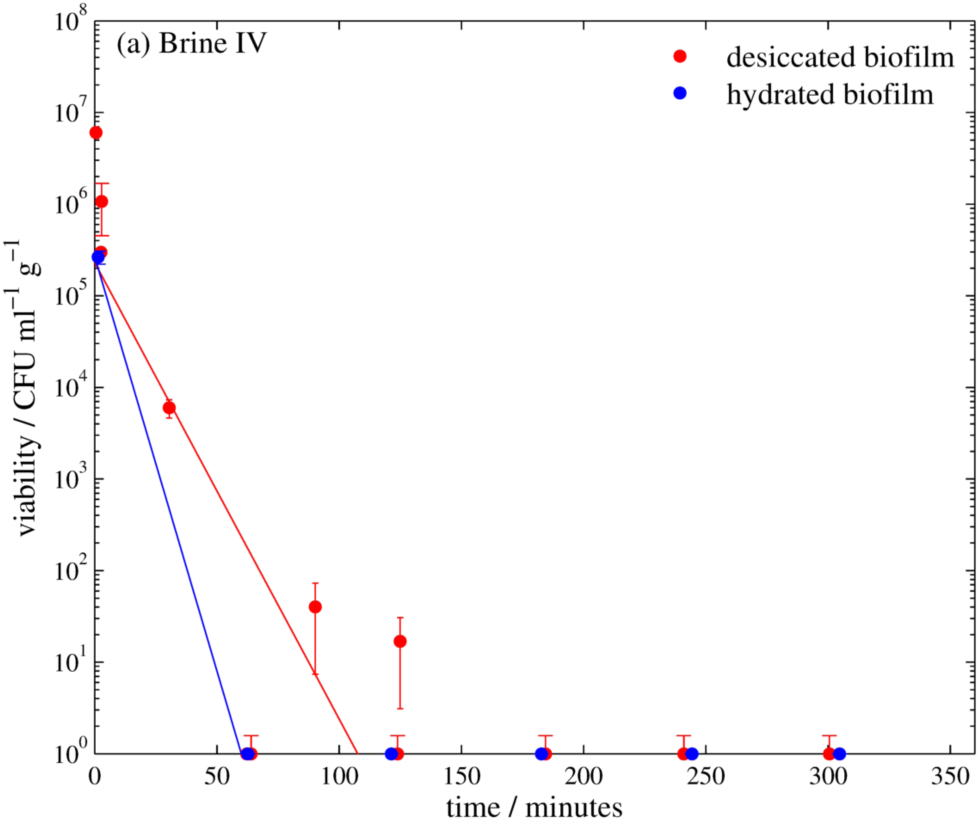
*Viability of S.* desiccabilis *cells in desiccated and hydrated biofilms after periods of immersion in Type IV brine, measured by counting colony forming units (CFU). Where no CFUs were countable, results are plotted at 10^0^. Each data point is a biological triplicate result, with standard error bars, and trend lines are plotted as linear regressions of data from several systematic replicates.*

Planktonic cultures of *S. desiccabilis* showed broadly similar responses to the 'weaker’ brines but far stronger responses to the ‘harsher’ brines compared to biofilm cultures. There was no decrease in viability over several hours of immersion in brines Ia, Ib and IIa. In brines IIb, IIIa, and IV there was a total loss in viability within less than 10 minutes (the shortest timepoint measured). In brine IIIb it was not possible to recover any planktonic cells into a pellet, suggesting they had lysed within 10 minutes.

To provide a more quantitative comparison of viability in the different brines, we calculated a linear regression for the results of each experiment and measured the gradient. Table 2 shows the gradient of these regression fits and the corresponding R^2^ value to assess the quality of the fits. A larger negative gradient implies a more rapid loss of viability, and a smaller R^2^ value suggests more variation in biofilm structure and growth between experimental replicates. The two instances of an R^2^ value of 1 are for experiments with only one measureable time point after the initial controls and so the fit is only for two points. These values are graphed in Figure 5. Brines IIb, IIIa, IIIb and IV show the most rapid loss of viability and there is a clear difference between desiccated and hydrated biofilms, with the latter losing viability ~2-10 orders of magnitude more rapidly when subjected to these brines.

**Table 2.** *Salts added to mix Mars analogue brines. Concentrations are in moles litre^−1^. Adapted from Fox-Powell et al. [15] and originally calculated from Tosca et al. [28].*

| Brine type | Ia | IIa | IIIa | Ib | IIb | IIIb | IV |
| --- | --- | --- | --- | --- | --- | --- | --- |
| NaHCO<sub>3</sub> | 0.126 |  |  |  |  |  |  |
| KHCO<sub>3</sub> | 0.028 | 0.041 |  | 2.237 |  |  |  |
| KCl | 0.022 | 0.020 | 0.075 | 3.776 | 1.033 | 1.142 |  |
| MgCl<sub>2</sub>·6H<sub>2</sub>O | 0.001 | 0.056 |  |  | 1.154 | 3.007 |  |
| NaCl |  | 0.154 | 0.189 | 1.266 | 2.265 | 1.036 |  |
| MgSO<sub>4</sub>·7H<sub>2</sub>O |  | 2.068 | 3.066 |  | 2.550 |  |  |
| FeSO<sub>4</sub>·7H<sub>2</sub>O |  |  | 1.225 |  |  | 2.313 |  |
| FeCl<sub>2</sub>·4H<sub>2</sub>O |  |  | 0.208 |  |  | 0.985 |  |
| HCl |  |  |  |  | 0.038 | 0.113 |  |
| Mg(ClO<sub>4</sub>)<sub>2</sub> |  |  |  |  |  |  | 3.097 |
| Water activity | 0.97 | 0.92 | 0.90 | 0.80 | 0.63 | 0.56 | 0.67 |
| pH | 8.2 | 6.2 | 1.8 | 8.3 | 2 | 0.6 | 5.5 |
| Ionic Strength | 0.18 | 8.5 | 12 | 5 | 11.6 | 13 | 9.29 |
| mol/liter |  |  |  |  |  |  |  |

**Figure 5.**
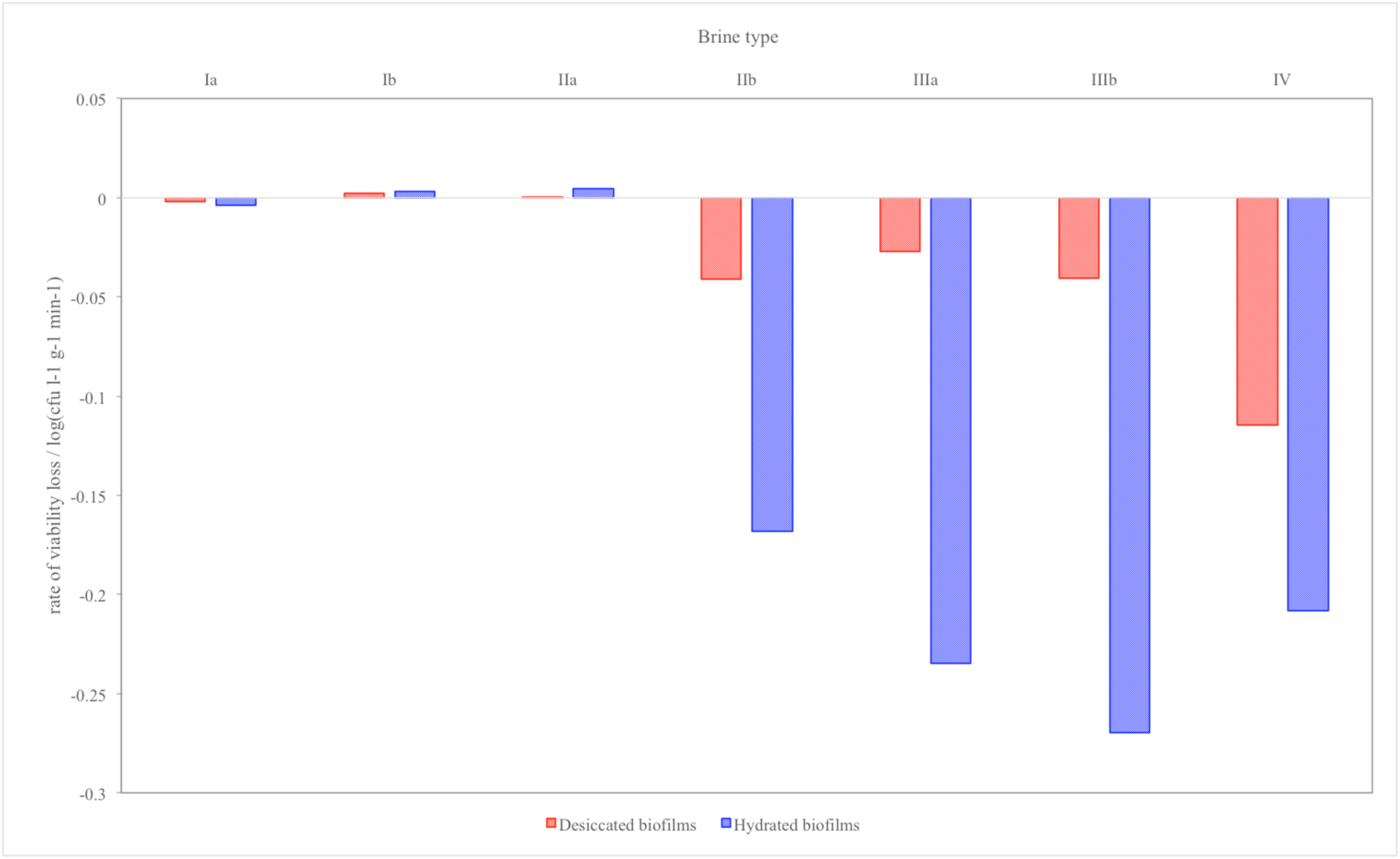
*Rate of viability loss of S.* desiccabilis *cells in desiccated and hydrated biofilms when immersed in Brines la to IV, measured from linear regressions of the curves shown in Figures 1–4.*

### 3.2 Repeated Brining Cycles

We also tested the viability of biofilms on scoria after repeated cycles of brining and desiccation, repeating the process used in Section 3.1. Viability was tested after two to five cycles of brine immersion. CFU counts after the extraction procedure were negative for all cycles tested.

### 3.3 Biomass assay

As Type I brines had little effect on the viability of cells in biofilms, these were not included in CV assays. Figure 6 shows the relative biomass present after a 24 hour incubation in the Type II, III and IV brines for both desiccated and hydrated biofilms. All of the hydrated biofilms showed a reduction in biomass after 24 hours, suggesting the biofilms were broken down by the brines. In the case of the desiccated biofilms, there was also a reduction in biomass after 24 hours in brines IIa and IV, but in brines IIb, IIIa and IIIb, the desiccated biofilms showed an increase in presumptive biomass over 24 hours, by up to 250%.

**Figure 6.**
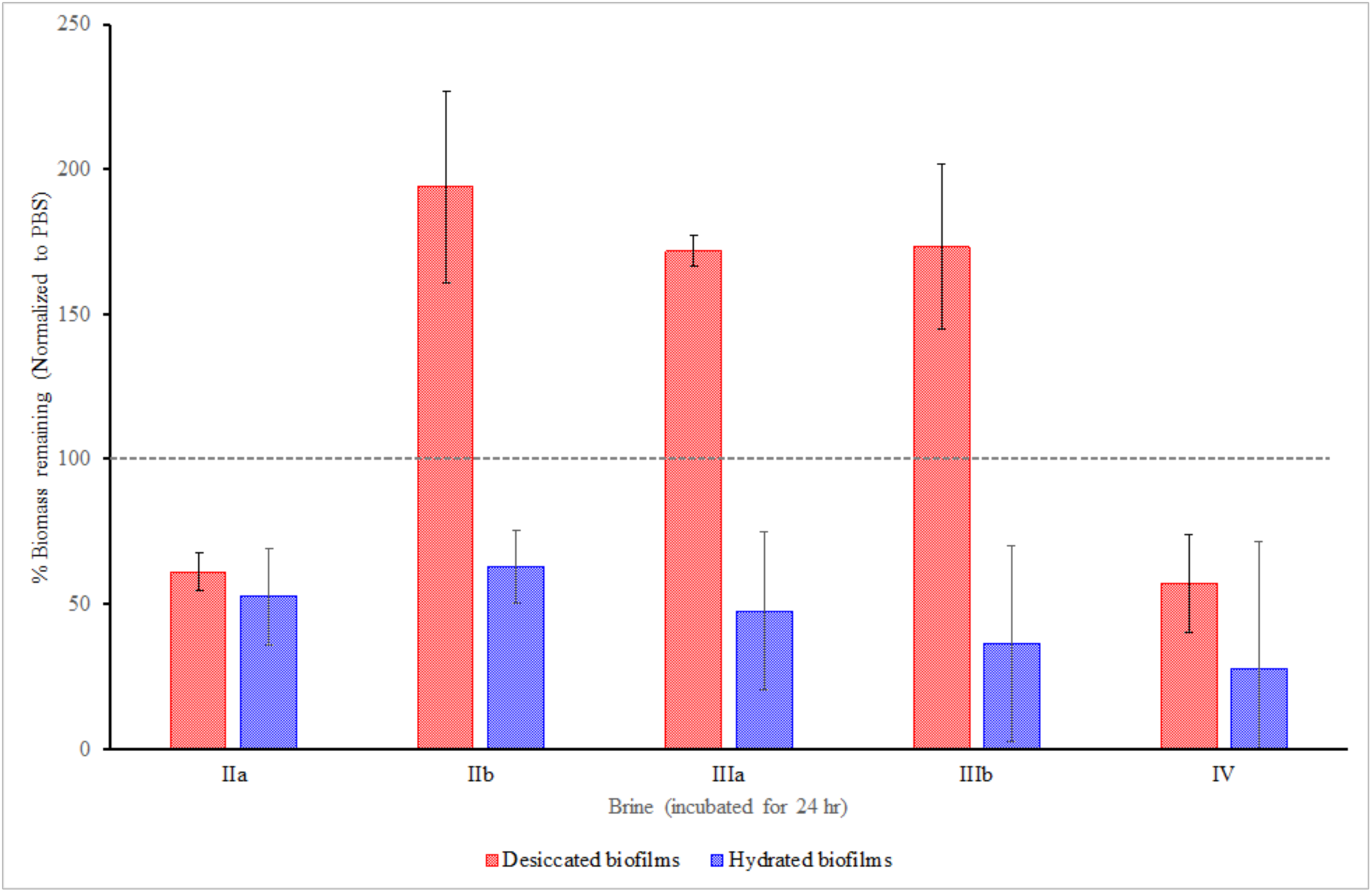
*Percentage biomass remaining after immersion in brines IIa to IV for 24 hours for desiccated and hydrated biofilms. Error bars are calculated from biological and systematic triplicates.*

To investigate this increase in biomass, we also performed a time series using the same experimental procedure, with brine IIb over another 24 hour period (Fig. 7). During incubation in the brine there was an initial reduction in biomass followed by an increase over several hours, which peaked at around 6 hours after exposure to brine. After this increase, the biomass began to reduce, from a peak of 200% to an average of around 100% after 24 hours.

**Figure 7.**
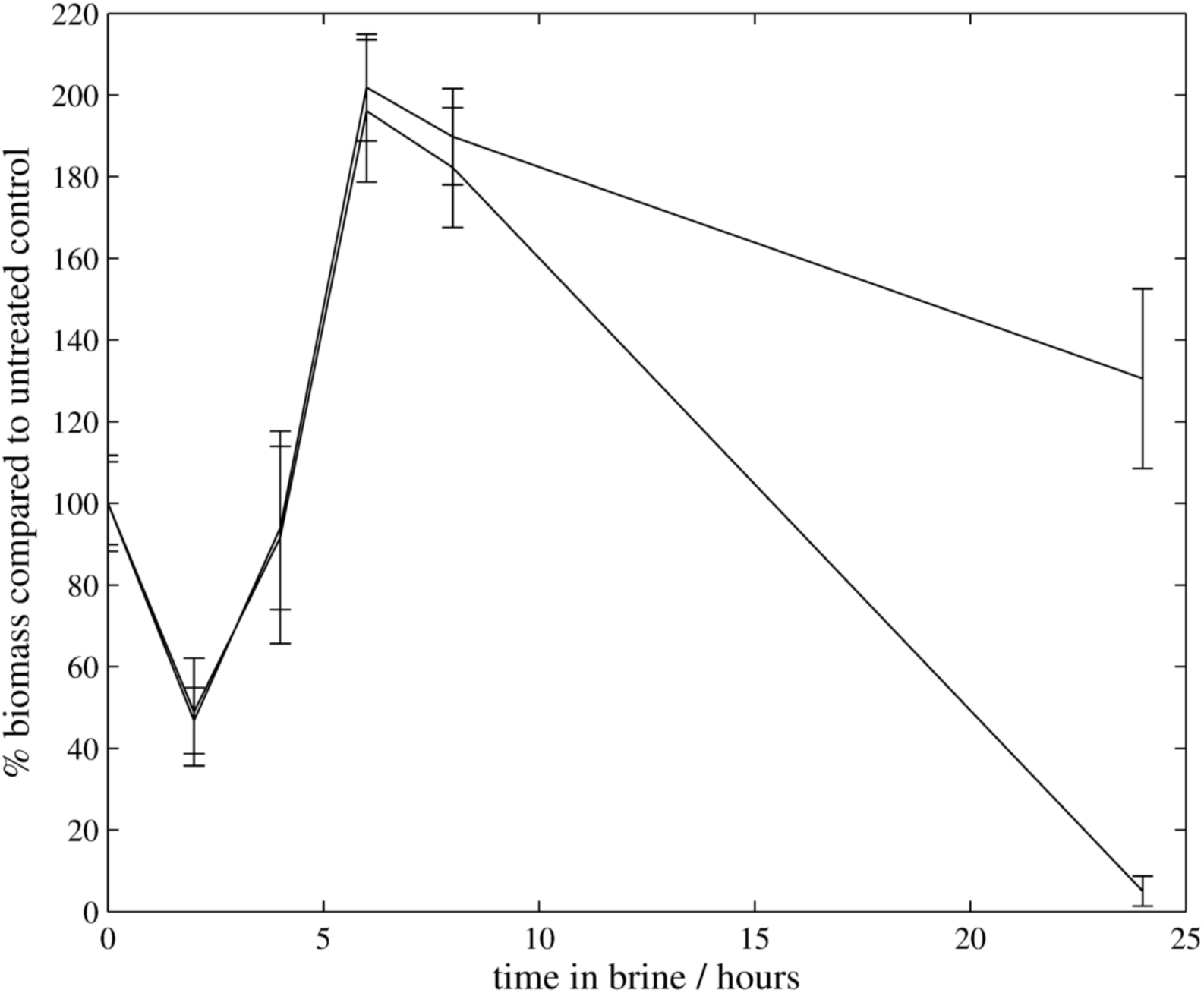
*Biomass remaining over time for desiccated biofilms in Brine IIb. Two experiments were performed.*

## 4 Discussion

Other planetary bodies in the solar system harbour aqueous environments that differ markedly in geochemical composition from aqueous environments on the Earth. These differences are a consequence of contrasting geological histories, which have produced the rocks and atmospheres that ultimately determine water chemistry.

In this study we sought to address the hypothesis that as on Earth, specific adaptations of microbial growth would influence the extent to which organisms could persist in extreme extraterrestrial aqueous environments. In particular, biofilms are known to provide resistance to chemical extremes [e.g. 4,7]. These data have implications for planetary protection that is concerned with whether terrestrial microorganisms could grow in extraterrestrial aqueous environments into which they are inadvertently introduced and, more speculatively, the adaptations that would be required for extraterrestrial life, if it exists, to grow in these alien geochemical extremes.

We found that combined physicochemical extremities of the brines had a significant effect on microbial survival. Generally, those brines with high water activity, low ionic strength and moderate pH show little effect on the viability of *S. desiccabilis* cells, whether grown in a planktonic state or in a biofilm.

Brines with low pH and high ionic strength generally had a large impact on viability, making planktonic cells unrecoverable in less than 10 minutes. Our data suggest that water activity is one factor that influences the ability of biofilms to survive brine exposure. For example, brine IIIa has a high water activity, low pH and high ionic strength and had a lesser impact on viability than brine IIb, which has a similar similar pH and ionic strength, but lower water activity. This observation is supported by the rapid loss of cell viability for planktonic cells and biofilms immersed in brine IIIb, which has the lowest water activity of the brines tested.

However, factors other than water activity also appear to play a role in loss of viability in biofilms. Brine IV caused a more rapid loss in viability than Brine IIb, although it has a higher water activity, more moderate pH and lower ionic strength. Brine IV is the only solution with perchlorate ions, which may suggest that these ions may be deleterious to cells, perhaps because of the oxidizing capability of perchlorate [20].

The resistance to loss of viability caused by the extreme brines is likely to be caused by a combination of physical protection and metabolic adaptation. Biofilms are known to provide a barrier between cells and the external hostile environments [3,21], in this case providing protection against the physicochemical extremes of the brine.

As shown by the more rapid reduction in viability measured with hydrated biofilms compared to desiccated biofilms, the physical protection offered by the biofilm is not the only factor that allows the cells to remain viable for longer in the brine. Desiccation adds yet another level of protection against extreme brine exposure.

The crystal violet assay suggested that when desiccated, biofilms showed an increase in presumptive biomass after several hours in the brine, a result not seen with hydrated biofilms and repeated consistently in desiccated biofilms. One biological explanation for these observations might be that desiccated biofilms are able to begin production of metabolites (such as membrane-stabilising disaccharides) or additional EPS during initial exposure to osmotic stress. Production of EPS is a known response to osmotic stress/desiccation [22]. Since matric (drying) water stress is equivalent to osmotic water stress from high concentration solutes [22], by drying the biofilms before brining, this process might act to ‘prepare’ the organisms by signalling the production of osmotic-resistance metabolites (potentially changing the composition of the biofilm), which might continue during exposure to brine prior to loss of viability. These observations merit further study.

Whilst our experiments show that under the most extreme brines, biofilms confer resistance to loss of viability, we were unable to recover viable cells after 5 hours in the harsher brines (Brines IIb, IIIa, IIIb and IV). We cannot rule out that other organisms or communities might be able to adapt to these extreme conditions, but our results do not show long-term survival. Furthermore, we tested the hypothesis that in the desiccated state, biofilms would enhance survival in transient aqueous systems, such as those proposed to occur in Martian RSLs. Under repeated cycles of desiccation and exposure to brine, which simulated the conditions that might occur in RSL, we found that viability was completely lost after the first cycle in all cases.

We have shown that biofilms formed by microorganisms offer protection against the extreme conditions of Mars-analogue brine solutions. Cells in biofilms desiccated before being immersed in brines remained viable for orders of magnitude longer than planktonic cells or hydrated biofilms under the same conditions. Some of these brines had low water activity, low pH and high ionic content that would otherwise make them uninhabitable [15], but *S. desiccabilis* biofilms retained viability in them for short periods of time, despite them quickly destroying *S. desiccabilis* cells in the planktonic state. Even though *S. desiccabilis* is not an extreme halophile, nor is it known to possess exceptional tolerance to osmotic shock, its biofilm forming capability provides resistance to high levels of dissolved salts. It was selected here for this ability to form biofilms and known desiccation tolerance. Halophilic biofilm-forming organisms that are exposed to geochemical regimes similar to those investigated here would likely possess even greater resistance.

These results have implications for the designation of RSL as ‘special regions’ for planetary protection purposes [23]. Given that terrestrial spacecraft almost certainly carry a bioload, some portion of which may be in biofilm form, even the transient aqueous environment suggested by RSL could present an environment that hardy terrestrial microbes could exploit or leave contaminating material in, especially if protected by a biofilm.

Finally, our results add to the expanding body of literature showing the protective properties of biofilms in extreme environments. Prior studies have shown biofilms’ capabilities to protect resistant cells against osmotic stress, but not with brines with properties as different or as extreme as those used in our study, such as concentrated sulfate brines. Although these brines are more relevant to extra-terrestrial environments, they are also relevant to some terrestrial locations such as the Basque lakes of British Columbia, which are sulfate-rich water bodies [24], or deep subsurface groundwater systems [25].

## 5 Materials and Methods

### 5.1 Strains and Growth conditions

This study examined viability and cell biomass in the organism *Sphingomonas desiccabilis* (*S. desiccabilis*), a Gram negative, rod-shaped, chemoheterotrophic, strictly aerobic bacterium originally isolated from a soil crust in the Colorado Plateau, USA [26]. Biofilms were cultured on pieces of basaltic scoria (highly vessiculated Icelandic lava (*Bar-Be-Quick Barbeque Products, Burnley, UK*) see Figure 8), chosen as an analogue for martian regolith and to provide a surface for the biofilm to form on. Overnight cultures of *S. desiccabilis* grown in R2A medium at 20°C [27], were normalised by cell density using OD_600_ measurements and inoculated onto ~5 mm pieces of scoria at a concentration of 50 μl in 4 ml of R2A medium and grown for one week at 20°C. Biofilm formation during this period was confirmed by visual inspection.

**Figure 8.**
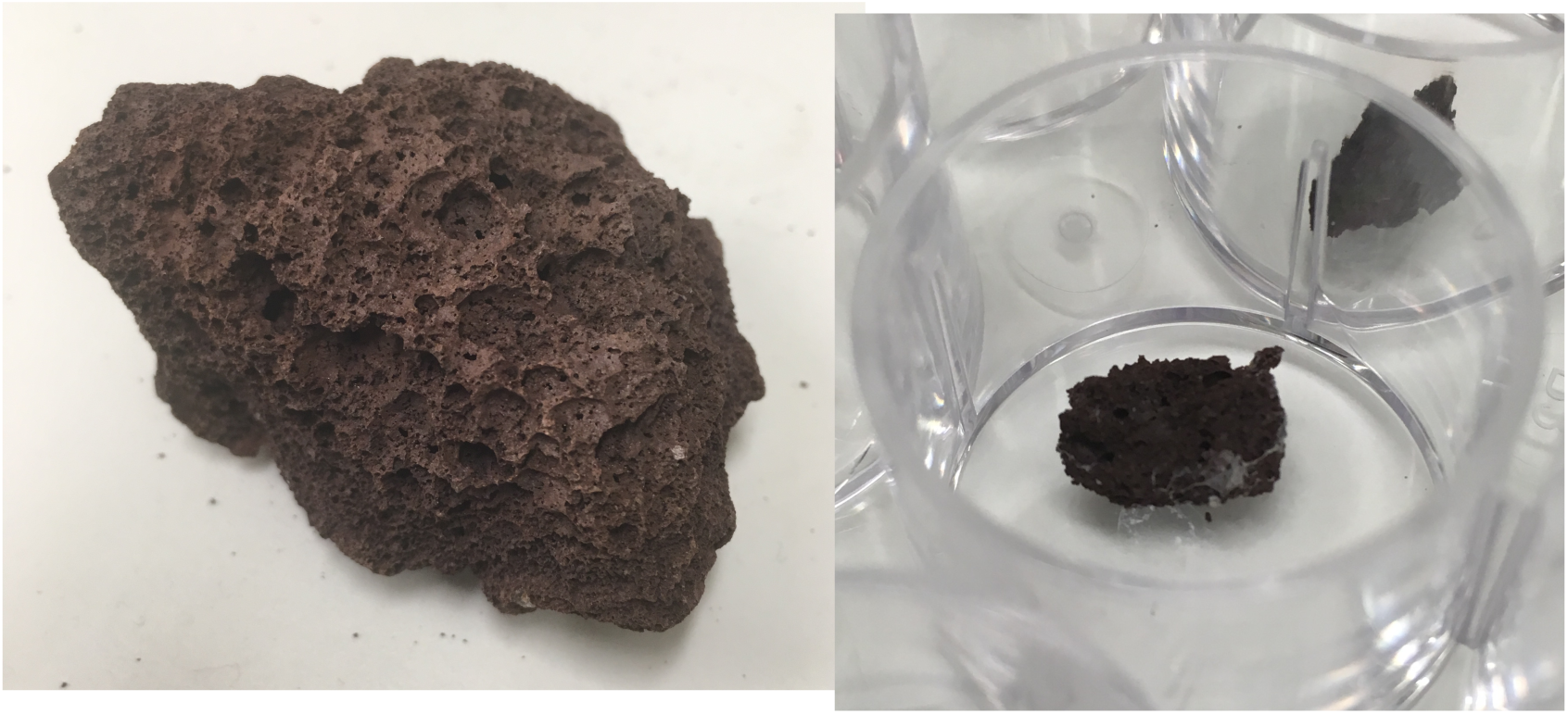
*Basaltic scoria used as a Mars analogue growth substrate a) larger piece and b) broken down to ~5 mm piece for use in well plates.*

After a week of growth, the supernatant was removed from the scoria. The biofilm-covered scoria pieces were then either a) dried in a laminar flow hood overnight or b) had their media replaced with PBS in order to halt growth but prevent desiccation. This allowed us to compare the response of desiccated biofilms to hydrated biofilms.

1-2 × 10^7^ cells planktonic cells were collected from 500 μl of the same normalised overnight cultures (incubated with shaking to reduce biofilm growth) for planktonic brine exposure experiments.

### 5.2 Exposure to brines

The biofilm inoculated scoria was then treated with Mars-analogue brines over a time course. The brines were synthesised based on computational reconstructions of putative martian brines by Tosca et al. [28], as used by Fox-Powell et al. [15], and include four types at two stages in their evaporation sequence (a and b). Type Ia and Ib are alkaline carbonate-chloride brines, which are analogous to the conditions in Gale Crater approximately 3.7 billion years ago [29] and the fluids that interacted with the Nakhla martian meteorite [30]. Type IIa and IIb are Mg-SO_4_-Cl dominated brines, characteristic of widespread Hesperian evaporite deposits such as those at Meridiani Planum [31]. Type IIIa and IIIb are similar in composition to Type II but contain higher levels of dissolved iron, and are extremely acidic.

In addition to the six Mars analogue brines, a magnesium perchlorate solution was also included (Type IV), given observations of magnesium perchlorate in the martian polar regolith by the Phoenix lander [32] and more recently associated with RSL following spectral observations [16]. This brine was a solution of magnesium perchlorate with a concentration to give a water activity matching that of brine IIb in order to investigate any differences caused by the chemical properties of the perchlorate salt. The molar recipes used to produce these brines are shown in Table 1, along with measurements of their physicochemical properties

**Table 1.** *Rate of viability loss of S.* desiccabilis *cells in desiccated and hydrated biofilms when immersed in Brines Ia to IV, measured from linear regressions of the curves shown in Figures 2–5.*

| Brine | Rate of viability loss / log(cfu l<sup>-1</sup> g<sup>-1</sup> min<sup>-1</sup>) |  |  |  |
| --- | --- | --- | --- | --- |
|  | Desiccated |  | Hydrated |  |
|  | Fit gradient | R<sup>2</sup> | Fit gradient | R<sup>2</sup> |
| Ia | -0.0018 | 0.098 | -0.0036 | 0.362 |
| Ib | 0.0022 | 0.095 | 0.0033 | 0.210 |
| IIa | 0.0002 | 0.083 | 0.0045 | 0.301 |
| IIb | -0.0411 | 0.770 | -0.1679 | 0.643 |
| IIIa | -0.0271 | 0.497 | -0.2346 | 1.000 |
| IIIb | -0.0407 | 0.358 | -0.2697 | 0.869 |
| IV | -0.1147 | 0.709 | -0.2081 | 1.000 |

Repeated brining was tested by removing the brines from the scoria after an hour. They were then dried in a sterile flow hood until there was no visible surface moisture on the scoria, and the process repeated.

### 5.3 Viability after brining

After being immersed in the brine, the scoria was washed twice in 1X PBS (Phosphate Buffered Saline) and then suspended in PBS. The suspension was vortexed briefly and sonicated for two minutes to extract cells from the biofilms. Planktonic cells that had been brined in the same manner were centrifuged at 8 g for 10 minutes, the brine removed and the cells re-suspended in 1 ml of PBS.

Cell suspensions were plated onto nutrient agar plates in serial dilution and the CFUs quantified after two days of growth at 20°C to give a measure of viability after immersion after a given amount of time in the different brines. Results were collected for each brine for both dried and hydrated biofilms in biological triplicate and systematic replicate at different time points.

### 5.4 Dettermination of Biomass

We performed a crystal violet (CV) staining to investigate how the brines affect the overall biomass of the biofilms. Biofilms were grown using the same method as described above and stained using a protocol adapted from Childers et al. [33]. The biofilm-coated scoria were incubated in the different brines, or PBS as a control for 24 hours, and then washed three times in PBS. Each scoria piece was then stained with 0.4% crystal violet solution for 15 minutes at room temperature, washed three times with PBS and destained for 15 minutes in 1 mL of 33% acetic acid solution. The optical density of the destain solution was used to quantify the total biomass of biofilm left on each piece of scoria after brining, with the PBS incubation forming the ‘zero brine’ control. The OD_570_ was measured by spectrophotometer with 33% acetic acid as the blank.

